# Achieving improved accuracy for imputation of ancient DNA

**DOI:** 10.1101/2022.04.26.489533

**Authors:** Kristiina Ausmees, Carl Nettelblad

## Abstract

Genotype imputation has the potential to increase the amount of information that can be gained from the often limited biological material available in ancient samples. As many widely used tools have been developed with modern data in mind, their design is not necessarily reflective of the requirements in studies of ancient DNA. Here, we investigate if an imputation method based on the full probabilistic Li and Stephens model of haplotype frequencies might be beneficial for the particular challenges posed by ancient data. We present an implementation called prophaser, and compare imputation performance to two alternative pipelines that have been used in the ancient DNA community based on the Beagle software. Considering empirical ancient data downsampled to lower coverages as well as present-day samples with artificially thinned genotypes, we show that the proposed method is advantageous at lower coverages, where it yields improved accuracy and ability to capture rare variation. The software prophaser is optimized for running in a massively parallel manner and achieved reasonable runtimes on the experiments performed when executed on a GPU.

## I. Introduction

The possibility of sequencing ancient human remains has allowed genetic analyses to be incorporated into studies of the history and evolution of our species. While methodological advances have led to an increase in the availability of ancient human genetic data, sample quality, quantity and cost still pose limitations for its analysis. Consequently, a significant amount of the available ancient DNA (aDNA) is sequenced at relatively low coverages of 1x or less, which leads some standard genomic and population genetic tools to be infeasible or unreliable to use [1].

One approach to extending the information content and making available a wider range of methods for such data is to impute genotypes at sites that are unobserved or otherwise not possible to call confidently. Imputation has been successfully applied to present-day data in a wide range of studies, including for conforming samples that have been genotyped on different arrays [2] as well as calling dense diploid genotypes from low- and medium-coverage data [3].

Genotype imputation is commonly performed by statistical models that take into account patterns of genetic variation in a panel of reference individuals with known haplotypes. The haplotype frequency model of [4] is the foundation for many commonly used imputation tools such as IMPUTE2 [5] and Beagle versions 4.1 and above [6]. Beagle 4.0 [7] uses a slightly different underlying model based on local haplotype clusters. Among more recent methods is GLIMPSE [8], which is designed for imputing low-coverage data and is based on a modified version of the Li and Stephens framework.

In the aDNA community, imputation has mainly been performed using the algorithm of Beagle 4.0 with genotype likelihoods as input [9]–[14]. In a recent study, an evaluation of different imputation pipelines on an empirical ancient sample was conducted [15]. They compared imputation performed in a single step based on genotype likelihoods, as implemented in Beagle 4.0 and GLIMPSE, to a two-step methodology in which either GLIMPSE or Beagle is used to determine sites which can be confidently called based on the likelihoods, after which imputation is performed by Beagle 5.0 based on the resulting genotypes. They found that the two-step methodology resulted in more refined posterior genotype probabilities that allowed for an improved balance between accuracy and number of retained sites when performing post-imputation filtering.

In general, methods for genotype imputation have been designed to target present-day data, with algorithm simplifications made in order to prioritize computational efficiency and applicability to increasingly large study samples and reference panels. While Beagle versions up to 4.0 allowed for imputation to be performed based on genotype likelihoods, later versions require genotypes as input, which may introduce errors for low-coverage data in which genotypes cannot be confidently called. Many methods also reduce the number of possible states induced by the phased reference, e.g. by the haplotype cluster model of Beagle 4.0, or by considering a subset of reference haplotypes based on some measure of similarity to the sample, as in IMPUTE2 and GLIMPSE. The latter two are also examples of models that decouple the phasing and imputation steps, iterating haplotype estimation based on sample data and subsequent haploid imputation. This is a strategy that can reduce the computational requirements of imputation, and has laid the foundation for pre-phasing approaches in which the two steps are completely separated and imputation requires phased input data. Also supported by IMPUTE2 and Beagle since versions 4.1, pre-phasing improves runtime for imputation with an often small trade-off in accuracy, although larger errors may result for populations where the haplotype phase is more difficult to estimate [16], [17].

These trends in priorities for the development of imputation methods do not necessarily reflect the requirements of aDNA studies, where the goal is often to extract as much information as possible from a few precious samples in order to perform population genetic analyses, rather than e.g. preparing tens or hundreds of thousands of samples for genome-wide association studies. Further, as low coverage and sequencing errors are prevalent in ancient data, and assumptions of the available phased reference panels being representative of the study samples may not hold, algorithm simplifications may incur larger trade-offs in accuracy than for present-day data.

In this paper, we evaluate the performance of the full probabilistic Li and Stephens framework for imputation of ancient data. We consider simulated low-coverage data based on present-day samples from the 1000 Genomes project as well as downsampled data from five empirical ancient individuals and compare performance to both a single- and a two-step imputation pipeline based on Beagle from [15]. Our implementation considers the full state space of possible haplotype phase configurations, which is integrated over to obtain posterior genotype probabilities. In order to obtain feasible runtimes on large reference sets, we have developed an implementation that is optimized for running on GPUs, which we make available in the GitHub repository https://github.com/scicompuu/prophaser.

## II. Methods

### A. Empirical data

Five ancient samples for which high-coverage genomes ranging from 19x to 57x are available were used for evaluation of imputation performance. These were sf12 from [18], Loschbour and LBK from [19], ne1 from [9] and ans17 from Fraser and Sánchez-Quinto *et al*. (in preparation). Gold standard genotypes to which imputed data was compared were obtained by the same pipeline as described in their respective publications, and further filtered to keep only sites that were confidently called. Low-coverage data for the evaluation samples was generated by downsampling reads to coverages ranging from 0.1x to 2x, after which genotype likelihoods were generated using a pipeline based on the The Genome Analysis Toolkit (GATK) v3.5.0 [20] tool UnifiedGenotyper. Unlike most previous studies of imputation of aDNA, we did not filter the imputation input to remove genotypes that could be affected by post-mortem damage, which was motivated by the findings in [15]. More information about the samples and data preparation procedure can be found in [14].

### B. Simulated data

Simulated low-coverage data was generated for 10 individuals from the CEU population of the 1000 Genomes phase 3 data set [3], filtered to keep only biallelic SNPs, by the following procedure.

For every site *j* with a biallelic SNP with called genotype *G_j_*:

- Decide whether or not to sample a read with probability *p*.
- If a read is not sampled, set the likelihood to 0.3333 for all 3 possible genotypes.
- If a read is sampled:

a. Sample one read from the distribution defined in Equation 1, which, for the biallelic SNP at location *j* that is sampled randomly *N_j_* times, describes the probability of sampling *A_j_* copies of the alternative allele A and *R_j_* copies of the reference allele R (so that *A_j_* + *R_j_* = *N_j_*). This assumes independently observed reads, and a constant sequencing error rate per base δ.

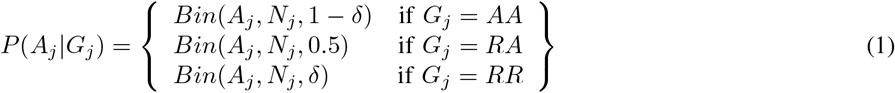
b. Let *X* denote the allele that was sampled, and *Y* the allele that was not sampled. Calculate likelihoods for the three genotypes according to Equation 2, which is based on the formula for genotype likelihoods from the GATK framework [20].

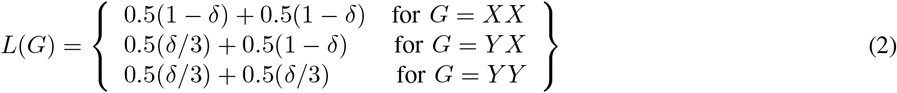

For different simulated coverage levels *c* ∈ [0, 1], the sampling probability *p* was selected so that the fraction of sampled sites would be approximately *c*. The called genotypes of the original 1000 Genomes data, *G_j_*, were used as gold standard genotypes to which imputed data was compared to assess accuracy. See the Supplementary Material for more information about the simulated data.

### C. Model and implementation

The method implemented in prophaser is based on a diploid version of the Hidden Markov Model (HMM) presented in [4]. We first outline this model, in which observed data corresponds to called genotypes, and later describe the implemented modification which instead considers genotype likelihoods.

For a set of *H* phased reference haplotypes at genomic positions *j* ∈ {1… *M*}, the observed states *G_j_* of the model correspond to the sample genotypes, and the hidden states *X_j_* = (*x*^1^, *x*^2^) consist of a pair of values that indicate which reference haplotype is used as a template for each chromosome at the position. The observation probabilities, shown in Table I, are based on the similarity between the observation and the genotype implied by the template haplotypes, and depend on an error parameter *ϵ_j_*. The transition probabilities of the HMM are shown in Equation 3. These depend on the number of switches in template haplotype for each chromosome, and are parameterized by *H* and *θ_j_*, reflecting the probability of historical recombination events between the two genomic positions. We refer to the Supplementary Material for more information about the model and its implementation.

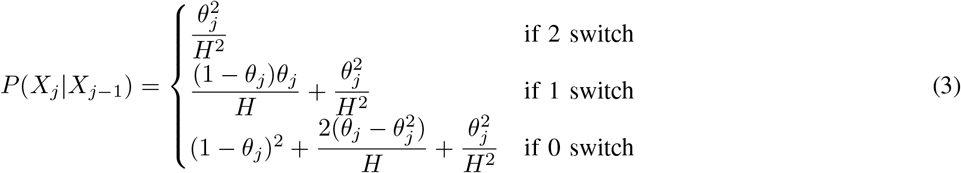

**TABLE I.** The observation probability *P*(*G_j_*|*X_j_*) of genotype *G_j_* at position *j* given the hidden state, with *G*(*X_j_*) denoting the genotype implied by the hidden state *X_j_*.

| | | $G(X_j)$ | | |
| --- | --- | --- | --- | --- |
| $G_j$ | | 0 | 1 | 2 |
| | 0 | $(1 - \epsilon_j)^2$ | $(1 - \epsilon_j)\epsilon_j$ | $\epsilon_j^2$ |
| | 1 | $2\epsilon_j(1 - \epsilon_j)$ | $(1 - \epsilon_j)^2 + \epsilon_j^2$ | $2\epsilon_j(1 - \epsilon_j)$ |
| | 2 | $\epsilon_j^2$ | $(1 - \epsilon_j)\epsilon_j$ | $(1 - \epsilon_j)^2$ |

The described model was used to define a HMM that incorporates the uncertainty of genotypes by having as observed data a vector of genotype likelihoods *L*(*g*) for each possible genotype *g*. For the case of biallelic SNPs, *g* ∈ {0, 1, 2} and the observation at position *j* is thus *Y_j_* = [*L*(0), *L*(1), *L*(2)]. Uniform likelihoods were assumed for unobserved sample sites. Integration over the unknown genotype yields the joint probability of hidden and observed states shown in Equation 4.

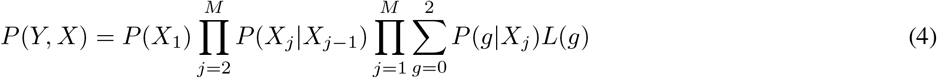

The forward-backward algorithm [21] was used to calculate posterior probabilities for the hidden states given the sequence of observed data *P*(*X_j_*|*Y*_1_,…, *Y_M_*) for all positions *j*. Phased haplotypes could then be obtained by selecting the state with the highest probability, and posterior genotype probabilities by integration over the hidden states.

Our implementation reuses the collapsing of transition probabilities involving switches between haplotypes outlined in [22]. This brings down the effective computational complexity to linear in the number of states (quadratic in the number of haplotypes). [22] also employed a checkpointing scheme, only keeping every *k* elements in the permanent forward-backward tables, reducing memory usage to be proportional to the square root of the number of variant positions. In our implementation, we extend the checkpointing scheme to two levels, rather than one. This makes memory usage proportional to the cubic root of the number of SNPs, with a trebling in the amount of computations relative to a non-checkpointing counterpart. The reductions in memory usage make the resulting algorithm more cache-friendly and the total workset small enough to fit in the local memory of a conventional GPU. Using OpenMP offloading pragmas, the same source code can be compiled for multicore CPUs and compliant GPUs.

### D. Experimental setup

Imputation performance was evaluated for three different methodologies: the proposed implementation prophaser as well as a single-step and a two-step pipeline based on Beagle. The single-step approach is the one that has mainly been used in aDNA studies previously, and consisted of using Beagle 4.0 to impute data using genotype likelihoods as input. The two-step methodology corresponds to the one that was recommended in [15] based on a comparison of single- and two-step pipelines based on Beagle and GLIMPSE. In this pipeline, Beagle 4.1 was used to estimate posterior genotype probabilities (GP) from the genotype likelihoods, to which a pre-imputation filter of maximum GP ≥ 0.99 was applied. The filtered genotypes were used as input to imputation, which was performed using Beagle 5.2. See the Supplementary Material for more details on the experimental setup as well as additional results.

Imputation was performed on chromosome 20 using a panel of phased reference haplotypes from the 1000 Genomes phase 3 data set and a genetic map derived from the HapMap2 [23] project. The phased reference data contained 2504 individuals and was filtered to include only biallelic SNPs. Experiments were performed on the simulated and empirical data sets separately. When imputing the simulated data, the 10 CEU individuals whose genotypes the sample data was derived from were removed from the reference set.

Imputation performance was assessed by genotype concordance, defined as the fraction of evaluated sites where the imputed genotype is the same as the gold standard genotype. We also report the number of SNPs evaluated in terms of raw numbers or the fraction they make up of the total amount of sites that are possible to compare, i.e. the number of sites that overlap the gold standard data and the non-filtered output from imputation. Where results are reported for different allele frequencies, the minor allele frequency (MAF) within the phased reference panel was used.

The Beagle pipelines were run on CPUs, with Beagle 4.0 allocated 2 cores of a compute server consisting of two 8-core Xeon E5-2660 processors, 218GB memory and Beagle versions 4.1 and 5.2 running on compute servers consisting of two 10-core Xeon E5-2630 V4 processors, 128 GB memory using 4 and 5 cores, respectively. The prophaser runs were performed on an NVIDIA A100 Tensor Core GPU with 40 GB on-board HBM2 VRAM.

## III. Results

For the results in this section, we only considered sites at which the gold standard genotype was heterozygous, as these are more challenging to impute and less subject to chance agreement to the reference majority allele. We also note that we did not see any tendencies of the methods to systematically mispredict homozygotes as heterozygote. A post-imputation filter requiring the GP value of the most probable imputed genotype to reach a certain threshold was also applied, with different thresholds used for the three imputation methodologies in order to obtain comparable numbers of evaluated SNPs. We found that the two-step Beagle methodology resulted in less nuanced posterior genotype probabilities, with many genotypes having the value 1.0, and consequently a relatively large amount of sites being retained. We therefore used a high threshold of 0.999 for this pipeline, and adjusted the other two accordingly with 0.9 for single-step Beagle and 0.8 for prophaser. We prioritized obtaining similar numbers of evaluated sites for the lower coverages in particular, as these are of main interest in this study and also the cases with largest differences in performance between the methods. We refer to the Supplementary Material for additional results considering all genotypes as well as different GP thresholds for the post-imputation filter.

For runtimes, we report the execution time per sample and imputation window, see the Supplementary Material for more information about how the different methodologies were executed. Using the computational resources described in the previous section, prophaser execution took 40 min. The runtimes of the Beagle software depended on coverage, and ranged between 2.2 and 7.12 hours for the single-step pipeline, and between 1 minute and 2.5 hours for the two-step pipeline.

### A. Empirical data

Figure 1 summarizes the performance of the three imputation methodologies for different coverages of the empirical data set. The single-step Beagle pipeline showed more variable outcomes for the different evaluation individuals, and was outperformed by the other approaches. The largest difference between the methods was seen for coverages below 0.5x, where prophaser had higher concordance while retaining a larger share of SNPs. Figure 2 shows the imputation performance over the allele frequency spectrum for coverages 0.1x, 0.5x and 2x. Again, the differences between methodologies were more pronounced for lower coverages, with prophaser outperforming the others consistently across the MAF spectrum at 0.1x coverage, but only showing discernible advantages at the lowest MAF values for 2x data.

**Fig. 1.**
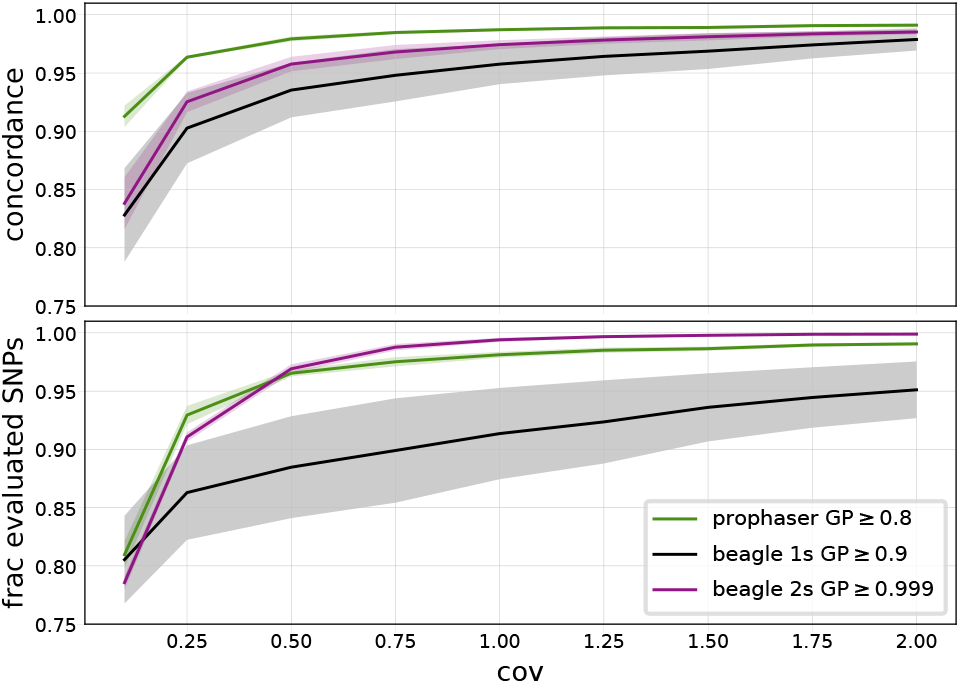
Genotype concordance (top) and fraction of evaluated SNPs (bottom) for heterozygote sites per coverage level, averaged over the five empirical evaluation individuals, with shaded regions displaying standard deviation. The legend indicates the GP thresholds used in the post-imputation filter for the different methodologies.

**Fig. 2.**
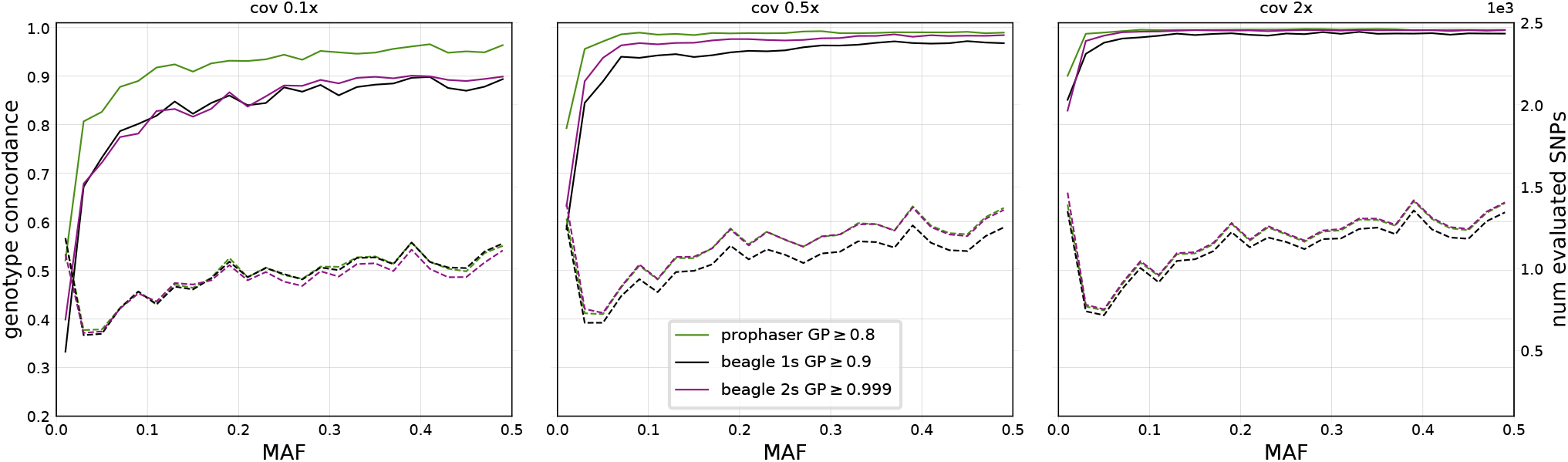
Genotype concordance (solid lines) and number of evaluated SNPs (dashed lines) for heterozygote sites, displayed over 25 MAF bins, averaged over the 5 empirical evaluation individuals, for three coverage levels. The three plots share both left and right y-axes, with axis ranges indicated on the respective figure edge. The legend indicates the GP thresholds used in the post-imputation filter.

### B. Simulated data

For the simulated data, the range of coverages considered contained lower values than that of the empirical data. Figure 3 shows that for coverages below 0.2x, the single-step Beagle methodology outperformed the two-step one, and had highest concordance overall for the lowest coverage considered of 0.01x. Inspection of the corresponding plot in Figure 4 shows that differences between prophaser and single-step Beagle varied over allele frequencies, where the latter had fewer sites retained at the lower end of the spectrum and a corresponding higher concordance, whereas the opposite holds at the higher end of the MAF range. Their relative performance may be subject to the particular post-imputation GP thresholds used, whereas the two-step Beagle pipeline had consistently lower concordance than the single-step, even when less sites were retained.

**Fig. 3.**
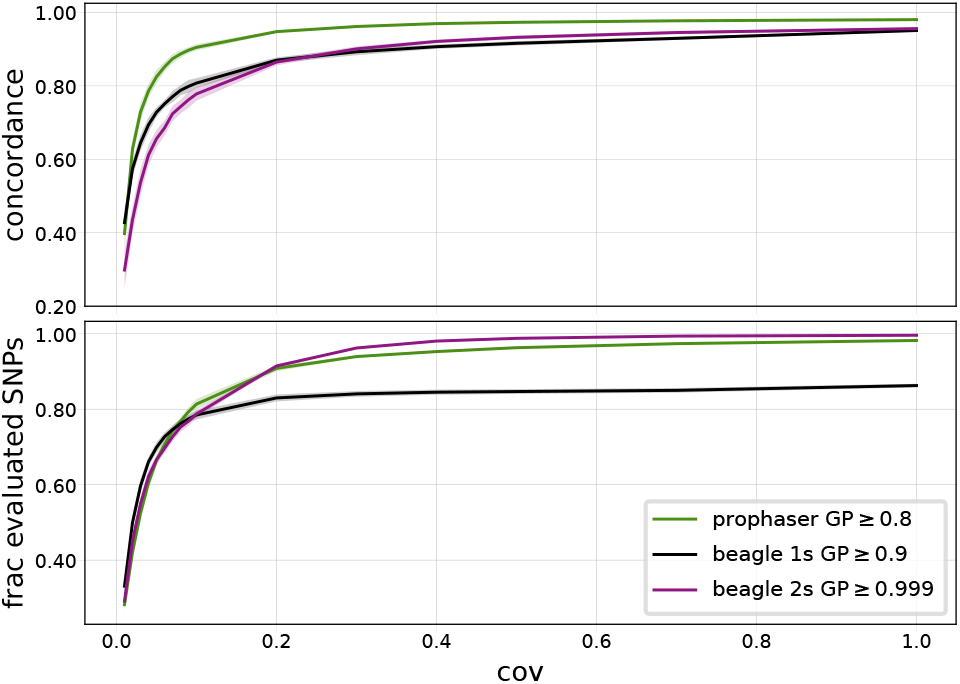
Genotype concordance (top) and fraction of evaluated SNPs (bottom) for heterozygote sites per coverage level, averaged over the 10 simulated evaluation individuals, with shaded regions displaying standard deviation. The legend indicates the GP thresholds used in the post-imputation filter for the different methodologies.

**Fig. 4.**
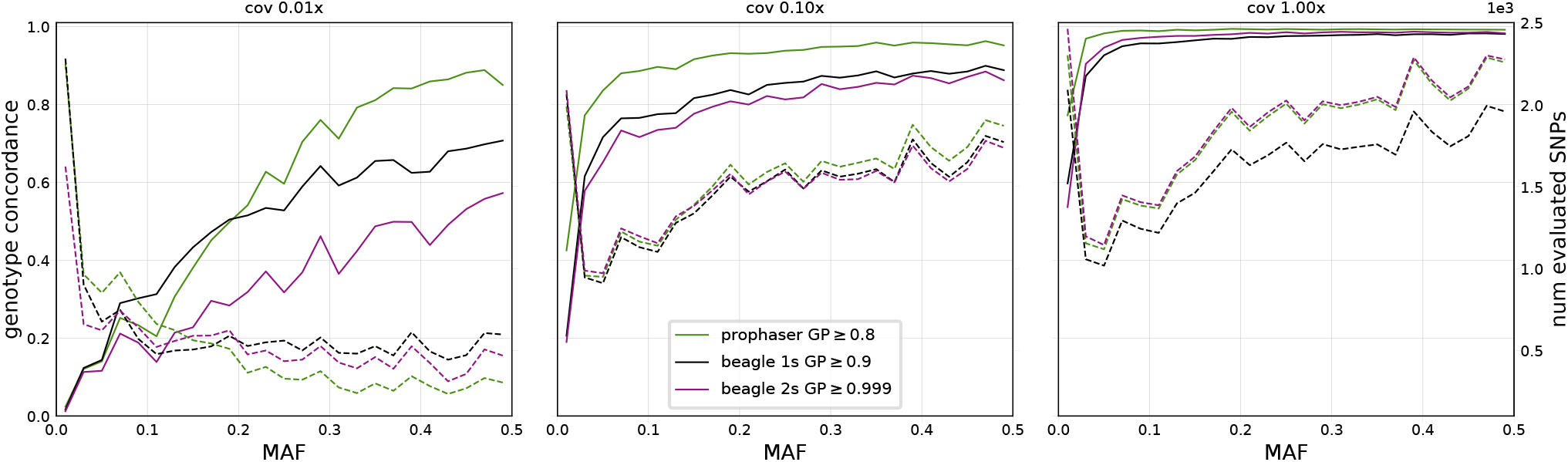
Genotype concordance (solid lines) and number of evaluated SNPs (dashed lines) for heterozygote sites, displayed over 25 MAF bins, averaged over the 10 simulated evaluation individuals, for three coverage levels. The three plots share both left and right y-axes, with axis ranges indicated on the respective figure edge. The legend indicates the GP thresholds used in the post-imputation filter.

For the higher coverages, the trends were largely similar to those observed for the empirical data set, with a main difference being increased performance for the single-step pipeline over the two-step one for coverage 0.1x, although this could also be driven by differences in number of sites retained by the filter. Another possible explanation for different patterns seen for the two data sets is that the haplotypes of the simulated samples are expected to have a higher degree of similarity to those in the reference panel as they are derived from the same population, in contrast to the ancient individuals which are temporally distinct from the present-day individuals in the panel.

## IV. Discussion and conclusion

We propose a method for imputation of missing genotypes that is based on the full probabilistic Li and Stephens framework for haplotype frequencies and show that it achieves higher heterozygote concordance than alternative methodologies based on the Beagle software on empirical ancient samples and simulated sparse genotypes. Our results suggest that the proposed method can provide reliable genotype calls over a larger part of the allele frequency spectrum and that it is particularly advantageous on low coverage data, and we therefore believe that it may be of use in studies of aDNA as a means to increase information content where high-coverage data is not available.

The presented implementation is optimized to run in a highly parallel manner, and we show that it achieves feasible runtimes on a realistic problem size when executed on a GPU. This design is in line with recent trends in the field of machine learning, where such processing architectures are seeing increased use due to applications like the training of deep artificial neural networks.

The underlying model lends itself to further customization for aDNA, and future work may focus on investigating the possibilities to include sample- or position-dependent adjustments to the HMM. This could include the incorporation of known patterns of errors in aDNA into the observation probabilities, or using estimates of divergence between sample and reference to adjust transition probabilities.

## Acknowledgements

The authors acknowledge the use of computational resources provided by Swedish National Infrastructure for Computing (SNIC) and associated centers under projects SNIC 2021/22-94, UPPMAX 2020/2-5 and SNIC 2017/7-162, as well as the use of the Berzelius resource funded by the Knut och Alice Wallenbergs Stiftelse (KAW) under project Berzelius-2021-30.

## Funding

This work has been supported by Formas grant 2017-00453.

## Notes

https://github.com/scicompuu/prophaser

### Competing Interest Statement

The authors have declared no competing interest.

