## Supplementary Material for "Achieving improved accuracy for imputation of ancient DNA"

#### 1 Methods

Unless otherwise stated, imputation was performed on windows of 60000 bp, with an overlap of 25000 bp. The methods were run separately on each window and the results subsequently merged. The tools `splitvcf` and `mergevcf` supplied along with the Beagle software were used for this, and the software `bcftools stats` v1.9 for assessing concordance. All imputation pipelines were run with a genetic map based on the HapMap2 map, interpolated to the sites in the imputation reference. The GitHub repository `https://github.com/scicompuu/prophaser` contains the map and scripts used to derive it.

##### 1.1 Simulated data

Simulated data was generated for coverage levels $c \in [1.0, 0.9, 0.8, 0.7, 0.6, 0.5, 0.4, 0.3, 0.2, 0.1, 0.09, 0.08, 0.07, 0.06, 0.05, 0.04, 0.03, 0.02, 0.01]$ using the sequencing error rate $\delta = 0.001$. The subsampling was done in an successive manner, so that independently for each individual, the sites that were sampled for a coverage level were a subset of the sites that were sampled for higher coverage levels. In the reported results, we omit coverages $0.9, 0.8$ and $0.6$ for brevity, as the differences between methods were relatively small for higher coverages.

##### 1.2 prophaser pipeline

In the underlying model for prophaser described in the main article text, the recombination parameter $\theta_j$ for a site $j$ is defined as $\theta_j = 1 - e^{\frac{-4N_e d_j}{H}}$, reflecting the fact that the transition rate depends on the genetic distance $d_j$ between loci $j-1$ and $j$ as well as the effective population size $N_e$. Genetic distances were obtained from the genetic maps used, and the value $N_e = 140000$ was used for all reported results. This value was reached by parameter evaluation on the simulated data set using different values in the range $N_e \in [25000, 140000]$. For the error parameter $\epsilon_j$, the value 0.001 was used for all sites for all reported results. This was selected to match the sequencing error rate $\delta = 0.001$ used to simulate the data. As these parameter values were found to give satisfactory results for the empirical data as well, no additional parameter evaluation was performed for this data set.

For each study sample, one iteration of the prophaser algorithm was executed, meaning that only the phased reference haplotypes were used in defining the haplotype model, current estimates of the haplotypes of other study samples were not used. This was done in order to be consistent with the other methods to get comparable results for the different imputation pipelines, see below.

#### 1.3 Beagle pipelines

Beagle 4.0 was run using the JVM memory settings `-Xss5m -Xmx32G` and program settings `impute-its=10 phase-its=10 window=60000 overlap=25000 gprobs=true`, with genotype likelihoods as input. The default value of 5 for number of phasing and imputation iterations was increased to 10, as the program documentation stated this would typically increase accuracy, which was also confirmed by parameter evaluation on data considered in this study.

Beagle 4.1 was run using the JVM memory settings `-Xss10m -Xmx32g` and program settings `modelscale=2.0 niterations=10 window=60000 overlap=25000 gprobs=true`, with genotype likelihoods as input. According to the program documentation, increasing the `modelscale` parameter from the default 0.8 can improve accuracy as well as runtime when estimating posterior genotype probabilities from genotype likelihoods. The number of iterations `niterations` was increased from the default 5 to 10 as the documentation states this gives higher accuracy, which was also found by evaluation on our data sets. We also evaluated setting the error parameter `err` to 0.001, but found that this mainly had an effect for very low coverages, where it led to slightly more sites kept and lower concordances than the default value of 0.0001, for the post-imputation thresholds considered here.

Beagle 4.0 and 4.1 were both run on each evaluation sample individually. This was in part because of their longer runtimes, which allowed for parallelization, but also because preliminary results suggested that this gave improved performance for lower coverages, which were of main interest for this particular study.

Beagle 5.2 was run using the JVM memory settings `-Xss10m -Xmx16g` and program settings `gp=true err=0.001`. As this version of Beagle had much lower execution times than the others, we did not run separate jobs to impute different windows of the data, but executed on the entire chromosome using default values for the `window` and `overlap` parameters. For each data set and coverage level, Beagle 5.2 was run on a joint file containing all evaluation samples as input.

We also evaluated different values of the parameters `ne` and `err` for Beagle 5.2. For the effective population size `ne` we evaluated values 10000 and 100000 in addition to the default value of 1000000. For the error parameter `err` we evaluated values 0.001 and 0.0001 in addition to the default value which is calculated based on the number of reference haplotypes, resulting in approximately 0.00002 for our experiments. We found that default `ne` and an `err` value of 0.001 resulted in more nuanced posterior genotype probabilities that yielded higher genotype concordances at levels of retained sites that were comparable to the other methods for the post-imputation thresholds considered here, particularly at the lower end of the coverage range.

### 2 Results

Sections 2.1 and 2.2 contain additional imputation results for the empirical and simulated data sets, respectively. For both sections, results are shown for two different post-imputation filtering setups. First, we show results when using a GP threshold of 0.99 for all pipelines, as this is a commonly used value that gives an indication of the qualities of the imputed data for the different methods. The second set of results is for the GP thresholds used in the main article text. These were adapted to yield comparable results between the different methodologies, and were 0.8 for prophaser, 0.9 for single-step Beagle and 0.999 for two-step Beagle. For the adapted GP thresholds, we also present results for each evaluation individual separately in Supplementary Tables S1-S4, attached in a separate file.

#### 2.1 Empirical data set

##### 2.1.1 Posterior genotype probability filter of 0.99

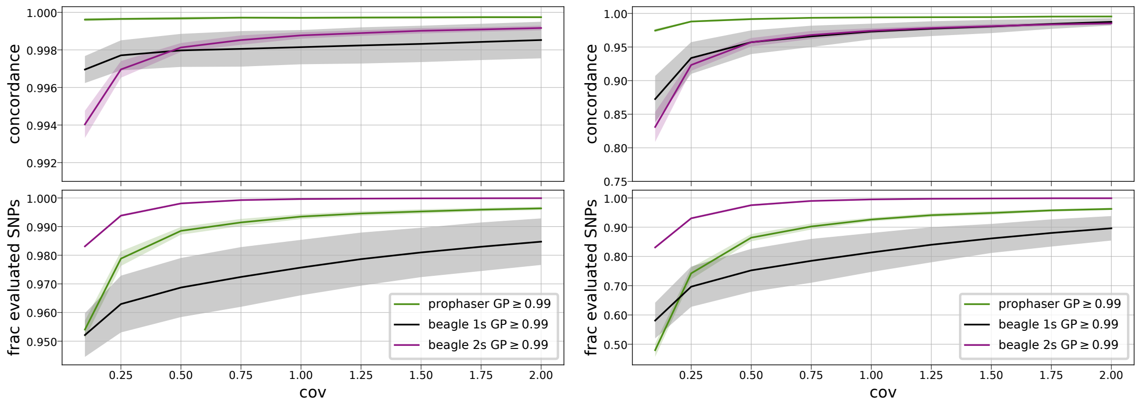


Figure 1: Genotype concordance (top) and fraction retained sites (bottom) per coverage level for the different imputation pipelines on the empirical data set. The left panel shows results for all genotypes, and the right panel for heterozygote genotypes. Results are averaged over the 5 samples of the empirical data set, with shaded regions showing standard deviation. Post-imputation filter thresholds are specified in the legend.

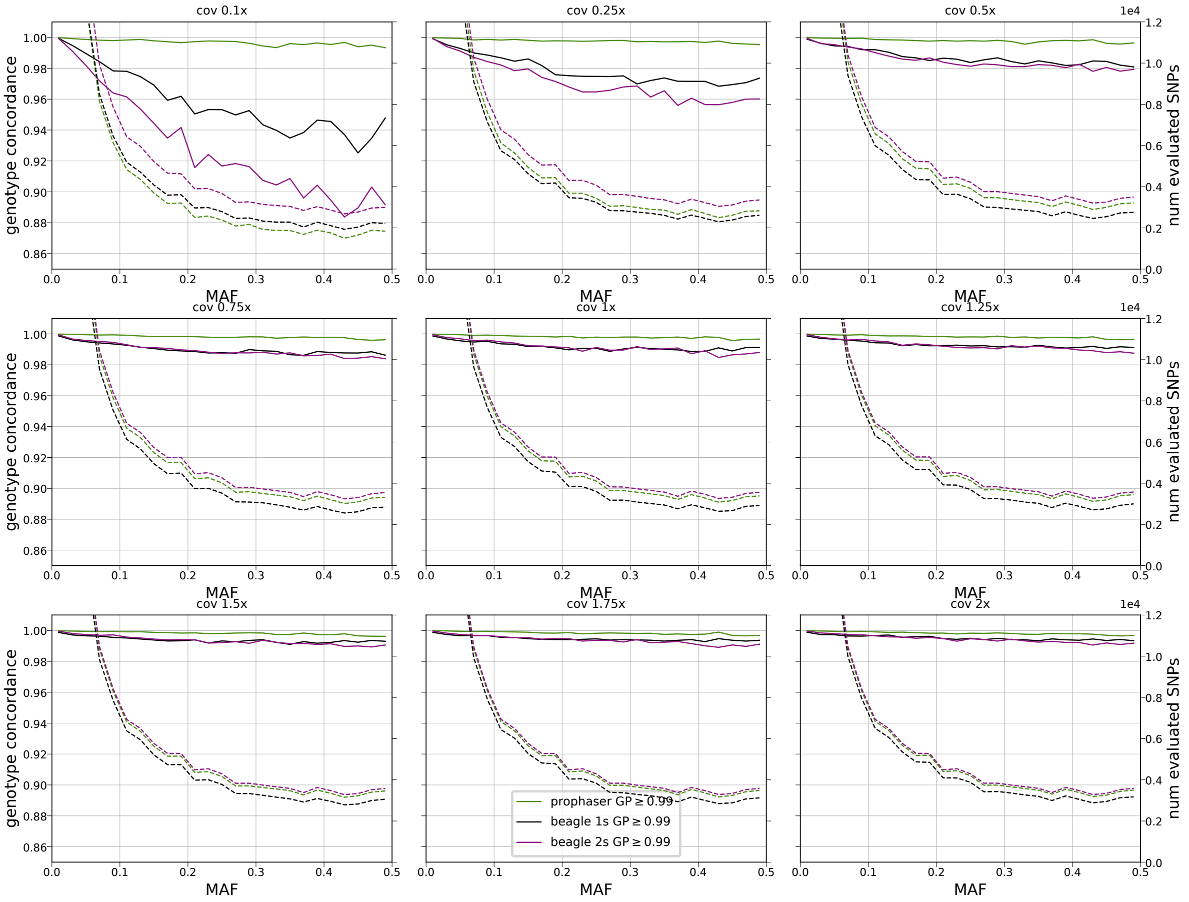


Figure 2: Genotype concordance per MAF (solid) and number retained sites (dashed), for all genotypes for different coverages, averaged over the 5 samples of the empirical data set, with post-imputation filter thresholds specified in legend.

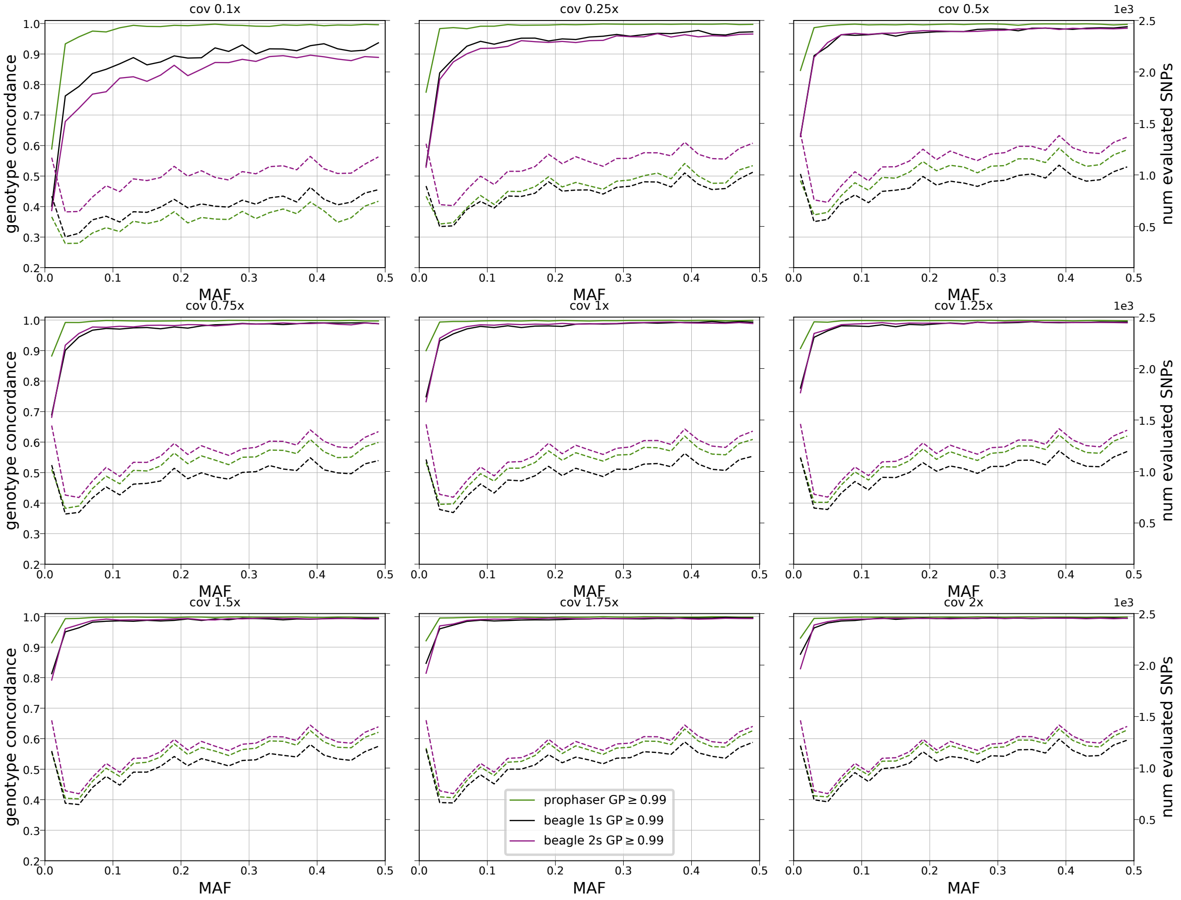


Figure 3: Genotype concordance per MAF (solid) and number retained sites (dashed), for heterozygote genotypes for different coverages, averaged over the 5 samples of the empirical data set, with post-imputation filter thresholds specified in legend.

##### 2.1.2 Adapted filter on posterior genotype probability

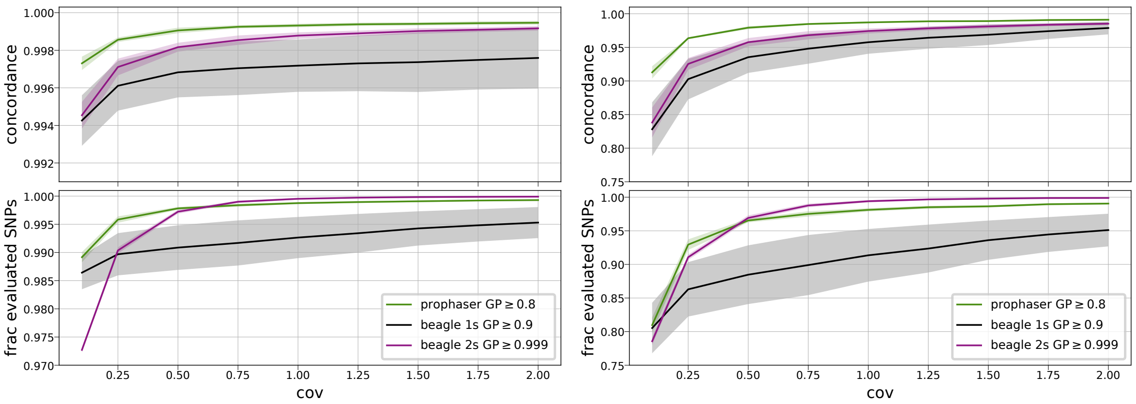


Figure 4: Genotype concordance (top) and fraction retained sites (bottom) per coverage level for the different imputation pipelines on the empirical data set. The left panel shows results for all genotypes, and the right panel for heterozygote genotypes. Results are averaged over the 5 samples of the empirical data set, with shaded regions showing standard deviation. Post-imputation filter thresholds are specified in the legend.

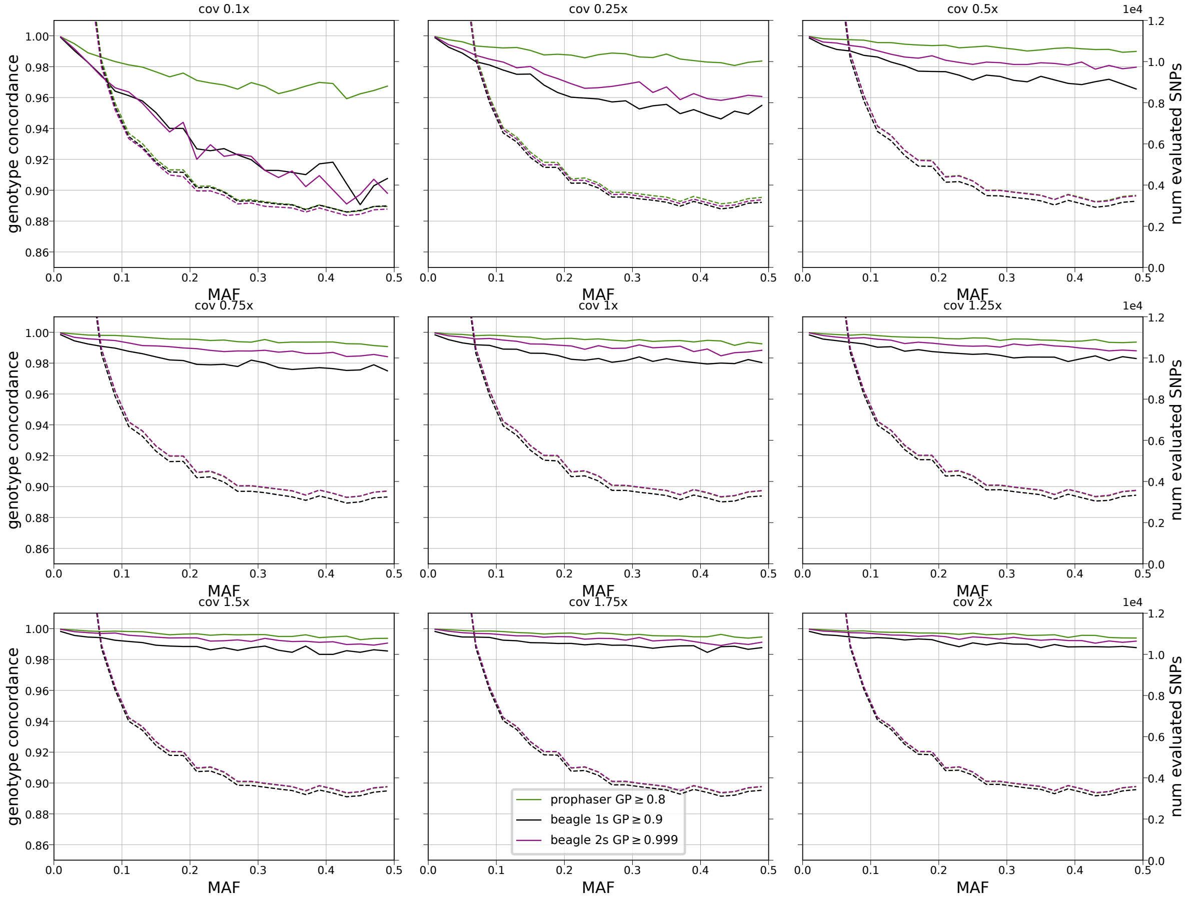


Figure 5: Genotype concordance per MAF (solid) and number retained sites (dashed), for all genotypes for different coverages, averaged over the 5 samples of the empirical data set, with post-imputation filter thresholds specified in legend.

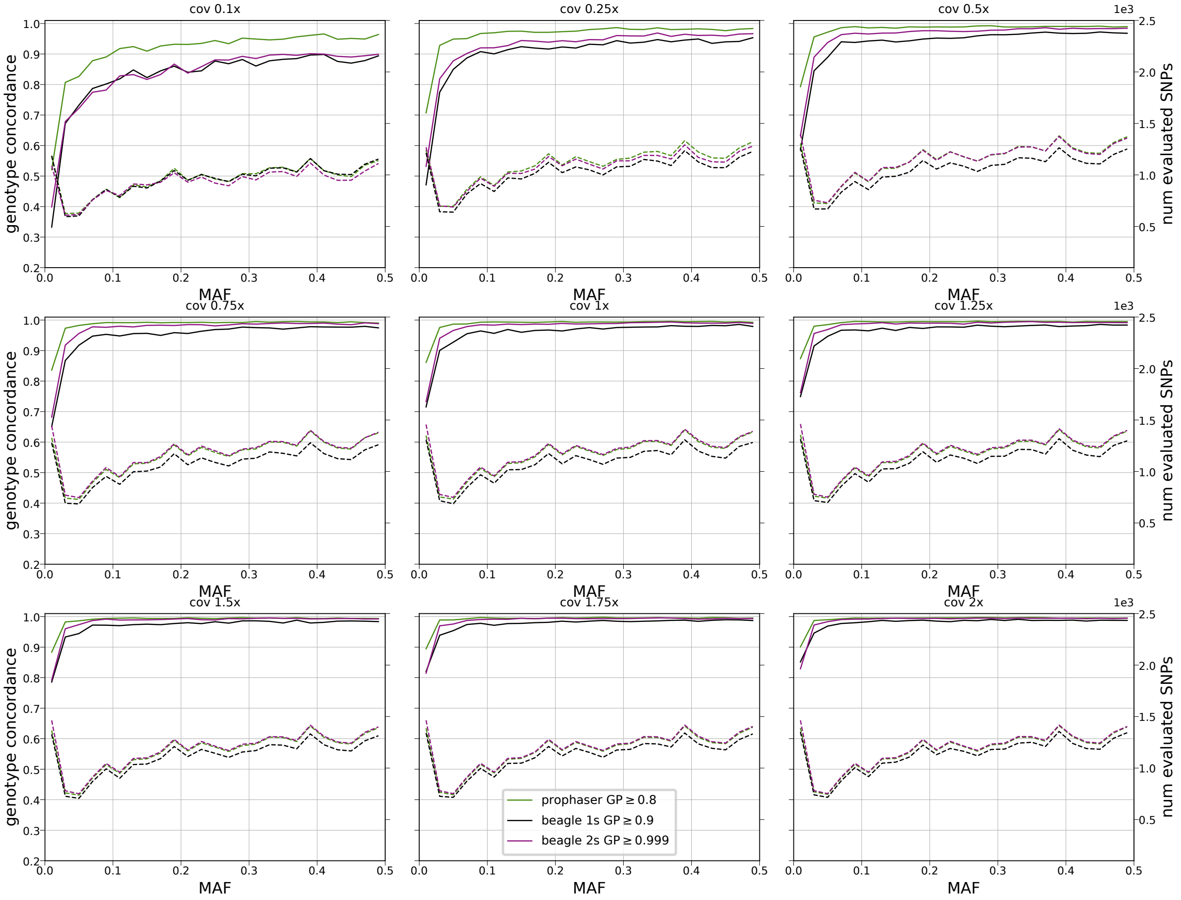


Figure 6: Genotype concordance per MAF (solid) and number retained sites (dashed), for heterozygote genotypes for different coverages, averaged over the 5 samples of the empirical data set, with post-imputation filter thresholds specified in legend.

#### 2.2 Simulated data set

##### 2.2.1 Posterior genotype probability filter of $0.99$

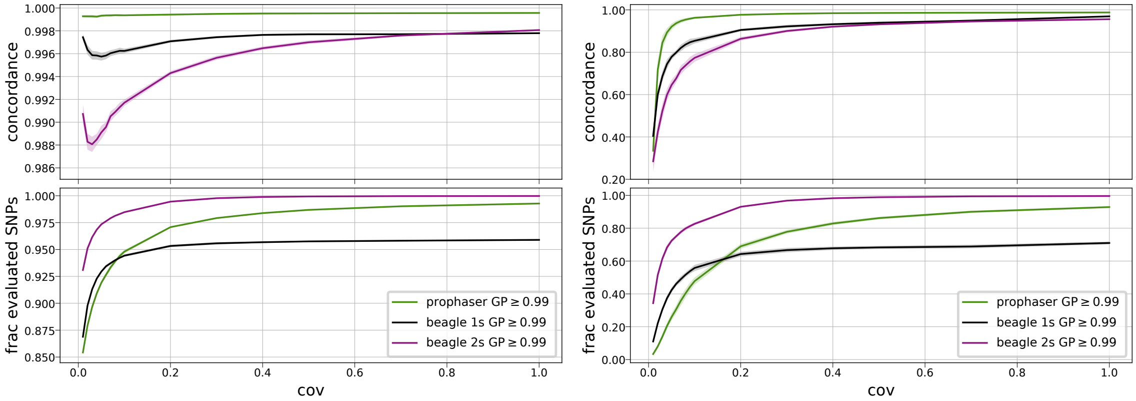


Figure 7: Genotype concordance (top) and fraction retained sites (bottom) per coverage level for the different imputation pipelines on the simulated data set. The left panel shows results for all genotypes, and the right panel for heterozygote genotypes. Results are averaged over the 10 samples of the simulated data set, with shaded regions showing standard deviation. Post-imputation filter thresholds are specified in the legend.

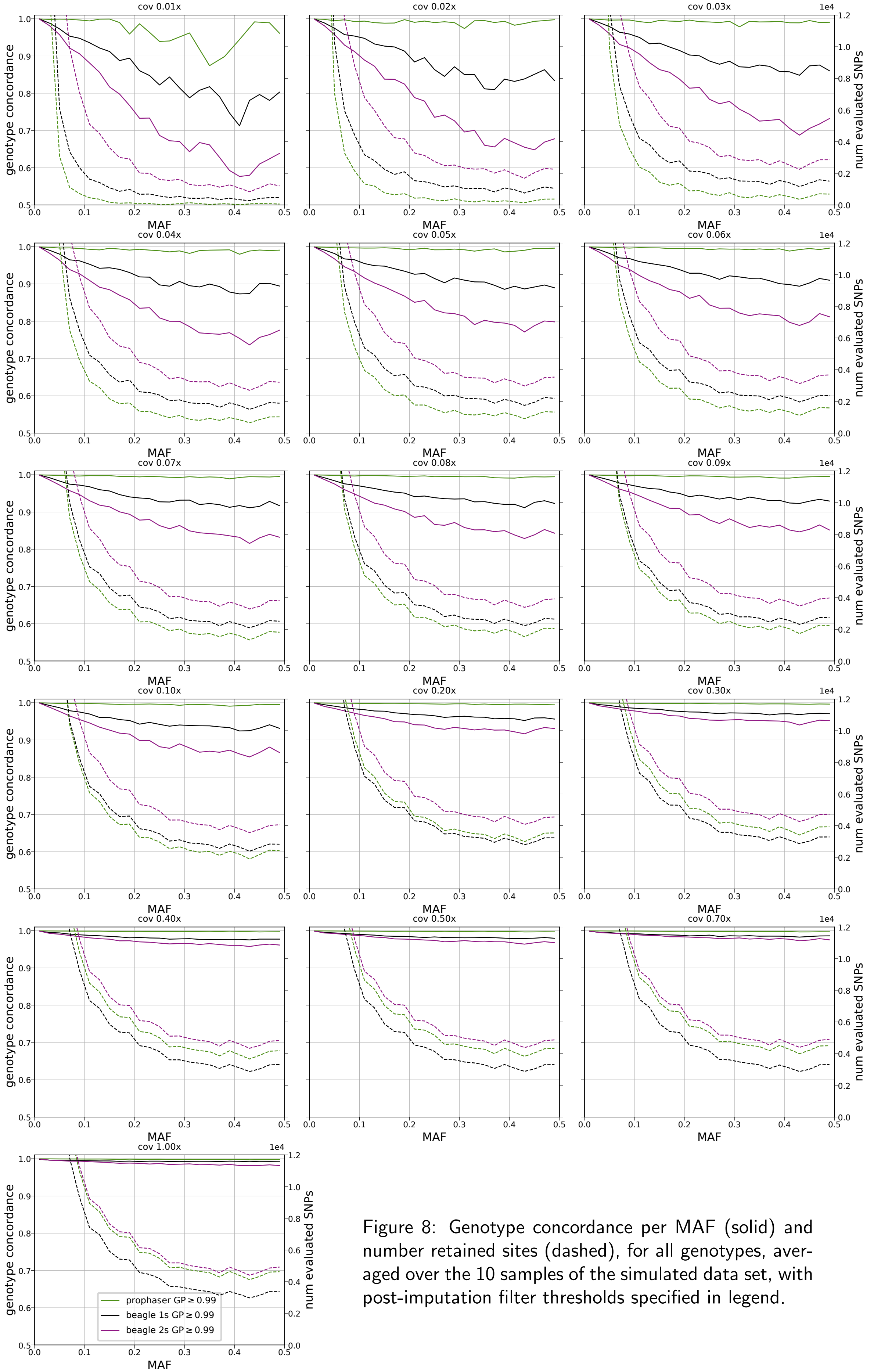


Figure 8: Genotype concordance per MAF (solid) and number retained sites (dashed), for all genotypes, averaged over the 10 samples of the simulated data set, with post-imputation filter thresholds specified in legend.

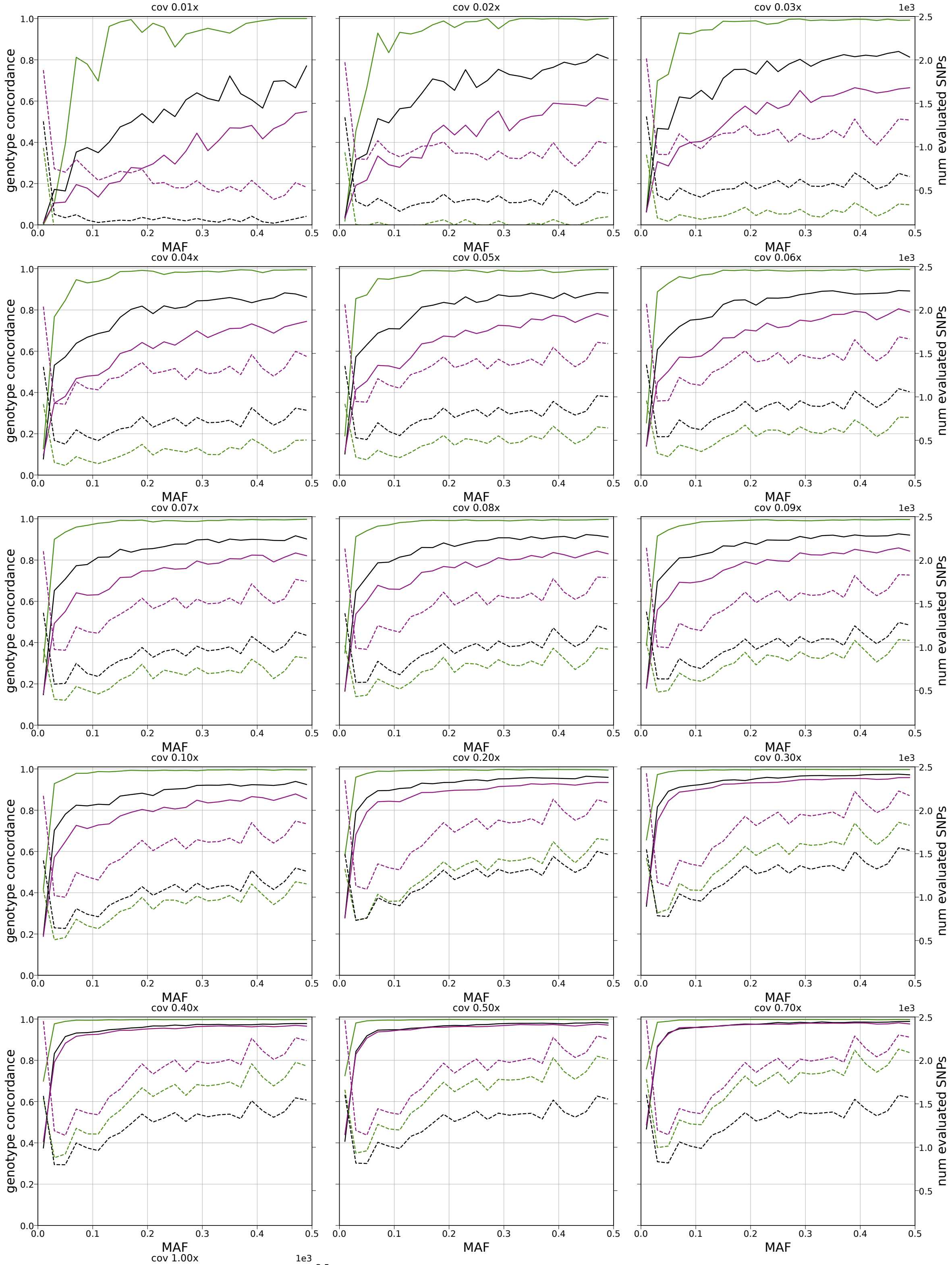


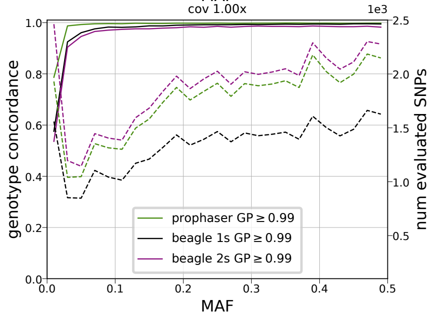


Figure 9: Genotype concordance per MAF (solid) and number retained sites (dashed), for heterozygote genotypes, averaged over the 10 samples of the simulated data set, with post-imputation filter thresholds specified in legend.

###### 2.2.2 Adapted filter on posterior genotype probability

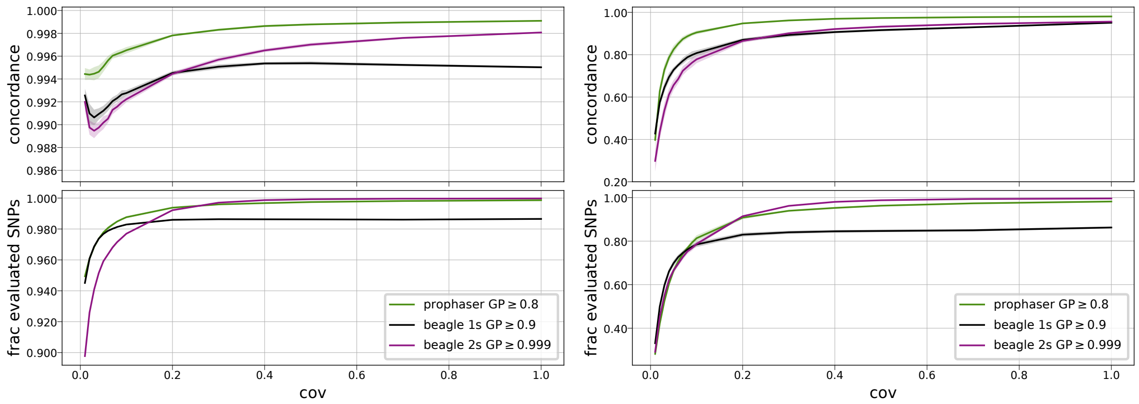


Figure 10: Genotype concordance (top) and fraction retained sites (bottom) per coverage level for the different imputation pipelines on the simulated data set. The left panel shows results for all genotypes, and the right panel for heterozygote genotypes. Results are averaged over the 10 samples of the simulated data set, with shaded regions showing standard deviation. Post-imputation filter thresholds are specified in the legend.

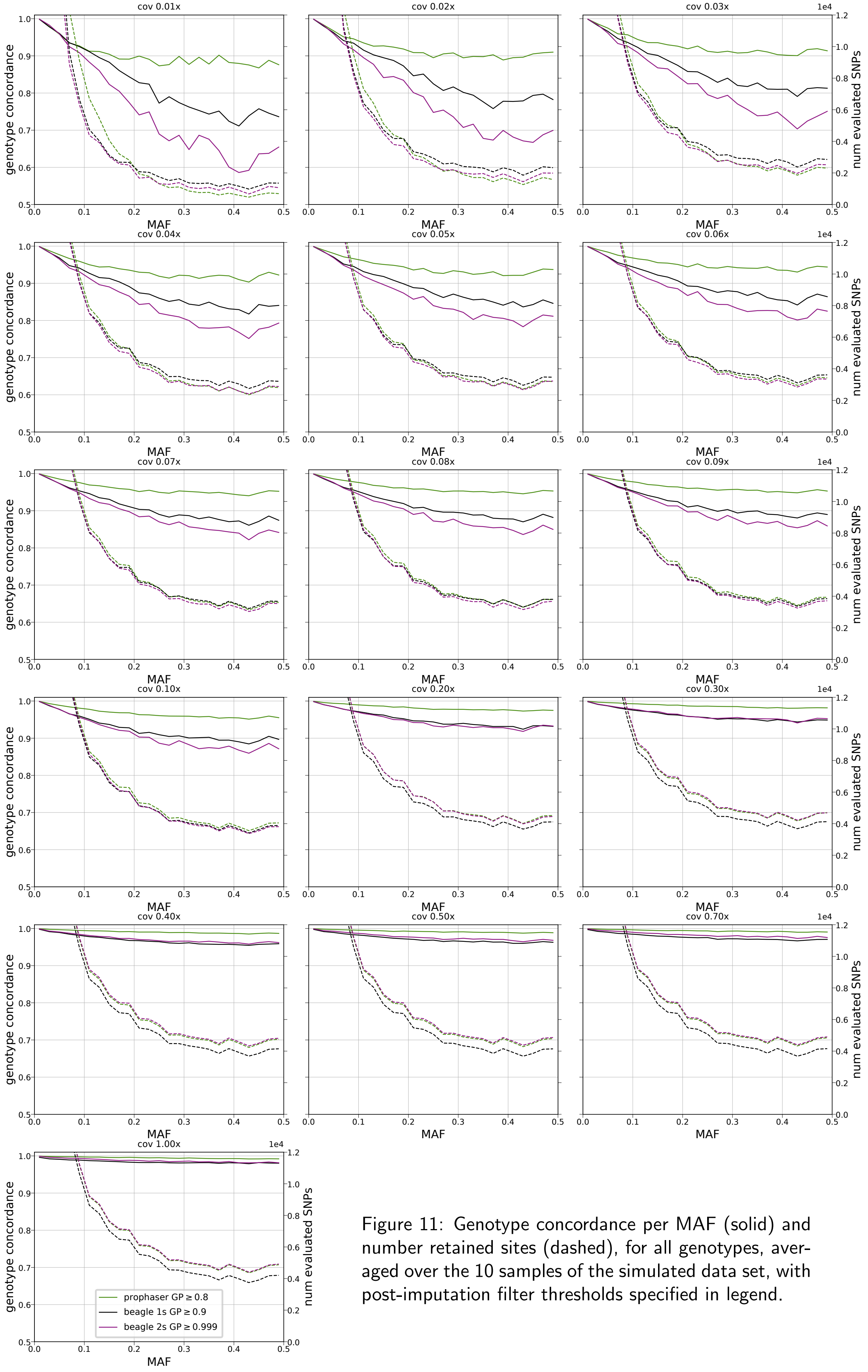


Figure 11: Genotype concordance per MAF (solid) and number retained sites (dashed), for all genotypes, averaged over the 10 samples of the simulated data set, with post-imputation filter thresholds specified in legend.

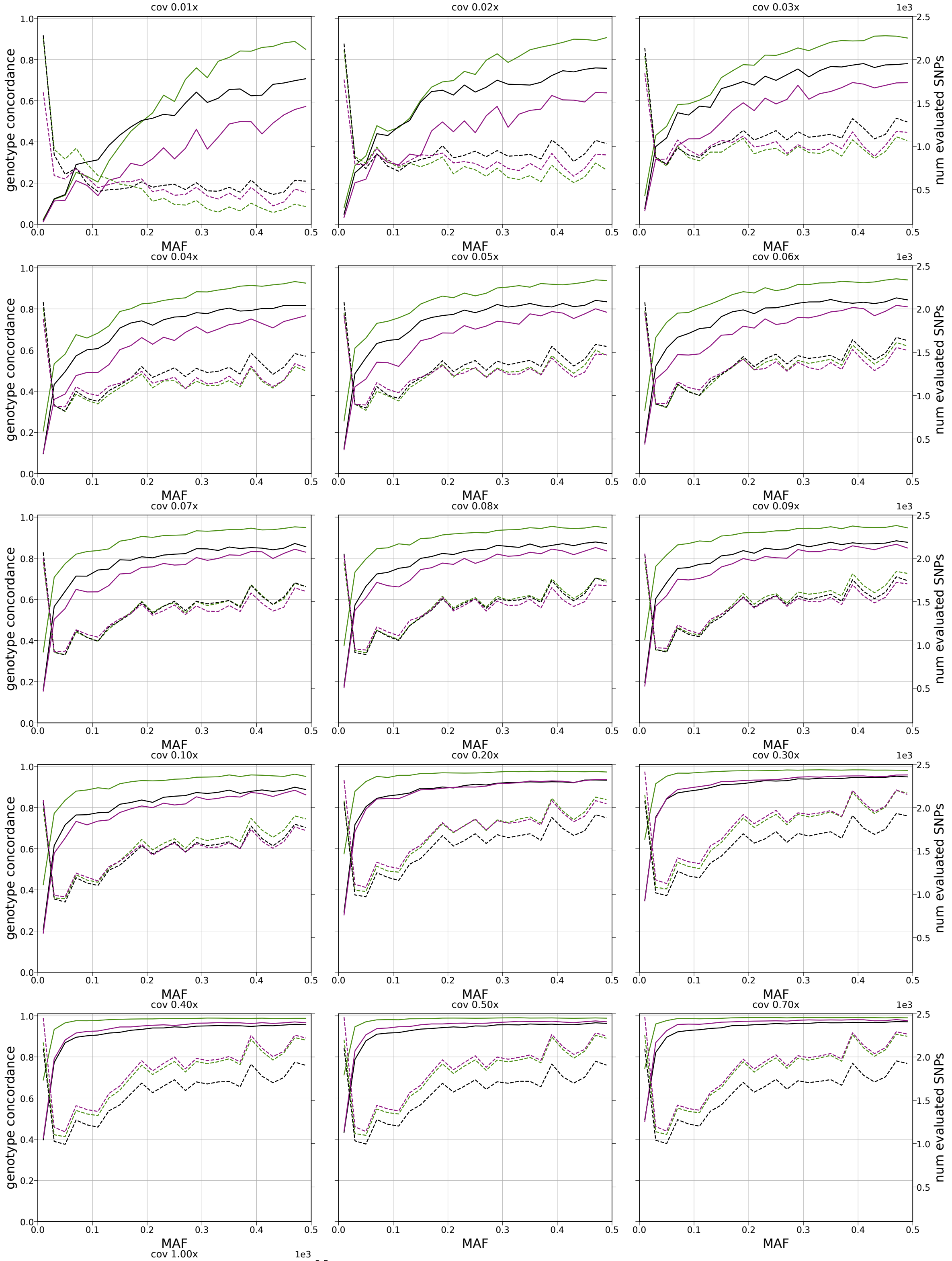


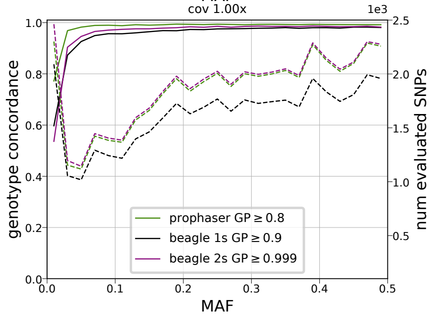


Figure 12: Genotype concordance per MAF (solid) and number retained sites (dashed), for heterozygote genotypes for different coverages, averaged over the 10 samples of the simulated data set, with post-imputation filter thresholds specified in legend.
